# Cell size and growth rate are modulated by TORC2-dependent signals

**DOI:** 10.1101/188698

**Authors:** Rafael Lucena, Maria Alcaide-Gavilán, Katherine Schubert, Maybo He, Matthew Domnauer, Catherine Marquer, Douglas R. Kellogg

## Abstract

The size of all cells, from bacteria to vertebrates, is proportional to the growth rate set by nutrient availability, but the underlying mechanisms are unknown. Here, we show that nutrients modulate TORC2 signaling, and that cell size is proportional to TORC2 signaling in budding yeast. The TORC2 network controls production of ceramide lipids, which play roles in signaling. We discovered that ceramide-dependent signals control both growth rate and cell size. Thus, cells that can not make ceramides fail to modulate their growth rate or size in response to changes in nutrients. PP2A associated with the Rts1 regulatory subunit (PP2A^Rts1^) is embedded in a feedback loop that controls TORC2 signaling and plays an important role in mechanisms that modulate TORC2 signaling in response to nutrients. Together, the data suggest a model in which growth rate and cell size are mechanistically linked by ceramide-dependent signals arising from the TORC2 network.

## Introduction

All cells control their size. From an evolutionary perspective, it would seem likely that cell size is controlled by highly conserved mechanisms, since size control would have been essential for the survival of the earliest cells. However, it has not yet been possible to define conserved mechanisms that can broadly explain cell size control in diverse cell types. Cell size control is relevant to cancer because severe defects in cell size are a nearly universal feature of cancer cells, yet nothing is known about the underlying causes.

The size of a cell ultimately reflects the outcome of mechanisms that control cell growth. One might therefore imagine that control of cell growth and size evolved together and are mechanistically linked. Indeed, there is evidence for a close relationship between cell growth and size. For example, in budding yeast, cell size at all cell cycle transitions is linearly proportional to the growth rate during the previous growth interval [1,2]. Moreover, the size of cells, from bacteria to vertebrates, is proportional to the growth rate set by nutrients [3-7]. Thus, cells growing in poor nutrients can be nearly half the size of cells growing in rich nutrients. A model that could explain the close relationship between growth rate and size is that the same signals that control growth rate also control the amount of growth that occurs during the cell cycle. Alternatively, cells could measure the rate of growth and set the amount of growth required for cell division accordingly. It is also possible that changes in cell size are an indirect consequence of changes in growth rate, such that there is no causal relationship between growth rate and size.

Distinguishing models for control of cell growth and size will require a better understanding of cell growth. Control of cell growth represents an extraordinary challenge. The central processes of growth - ribosome biogenesis and membrane expansion - must be coordinated with each other, and the rates of each process must be matched to the availability of building blocks and energy derived from nutrients. In budding yeast, the transcription of over a thousand genes is proportional to growth rate, which suggests that global top-down signals ensure that the rate of growth is matched to nutrient availability [8].

A conserved signaling network is thought to control cell growth in all eukaryotic cells. At the heart of the network are the Tor kinases, which assemble distinct multiprotein kinase complexes called TORC1 and TORC2 [9-12]. The TORC1/2 kinase complexes control members of the AGC kinase family, which play essential roles in controlling the diverse events of growth [13-15]. The AGC kinases are also controlled by phosphoinositide-dependent kinase 1 (PDK1) [16,17]. TORC1/2 and PDK1 phosphorylate distinct sites on AGC kinases that promote activation.

Little is known about how the diverse events of cell growth are coordinated with each other, or how growth rate and cell size are influenced by nutrients. In previous work, we discovered that PP2A^Rts1^, a form of PP2A associated with the Rts1 regulatory subunit, is required for nutrient modulation of cell size [18]. In addition, inactivation of PP2A^Rts1^ abolishes the linear relationship between cell size and growth rate [2]. Together, these observations suggest that PP2A^Rts1^ could play an important role in mechanisms that control cell growth and size. Here, we investigated the mechanisms by which PP2A^Rts1^ influences cell growth and size in budding yeast.

## Results

### PP2A^Rts1^ restrains the activity of Mss4, a conserved PI(4)P kinase

To investigate how PP2A^Rts1^ influences cell growth and size, we first searched for proteins controlled by PP2A^Rts1^. In previous work, we used proteome-wide mass spectrometry to identify proteins that are hyperphosphorylated in *rts1*Δ cells, which would suggest that they are direct or indirect targets of PP2A^Rts1^ [19]. Mss4, the budding yeast homolog of phosphatidylinositol-4-phosphate 5-kinase, emerged from this analysis as a strong candidate. The mass spectrometry identified 6 high-confidence sites on Mss4 that undergo hyperphosphorylation in *rts1*Δ cells.

An important function of Mss4 is to promote TORC2-dependent phosphorylation of Ypk1 and Ypk2, which are partially redundant AGC kinase paralogs that are homologs of vertebrate SGK kinases [17,20]. Mss4 produces PI(4,5)P_2_, which recruits the proteins Slm1 and Slm2 to the plasma membrane [21]. The Slm 1/2 proteins, in turn, bind both TORC2 and Ypk1/2, serving as a scaffold to drive phosphorylation of Ypk1/2 by TORC2 [20,21]. Full activation of Ypk1/2 requires further phosphorylation by Pkh1 and Pkh2, another pair of partially redundant AGC kinase paralogs that are the budding yeast homologs of vertebrate PDK1 (**Figure 1A**) [17].

**Figure 1:**
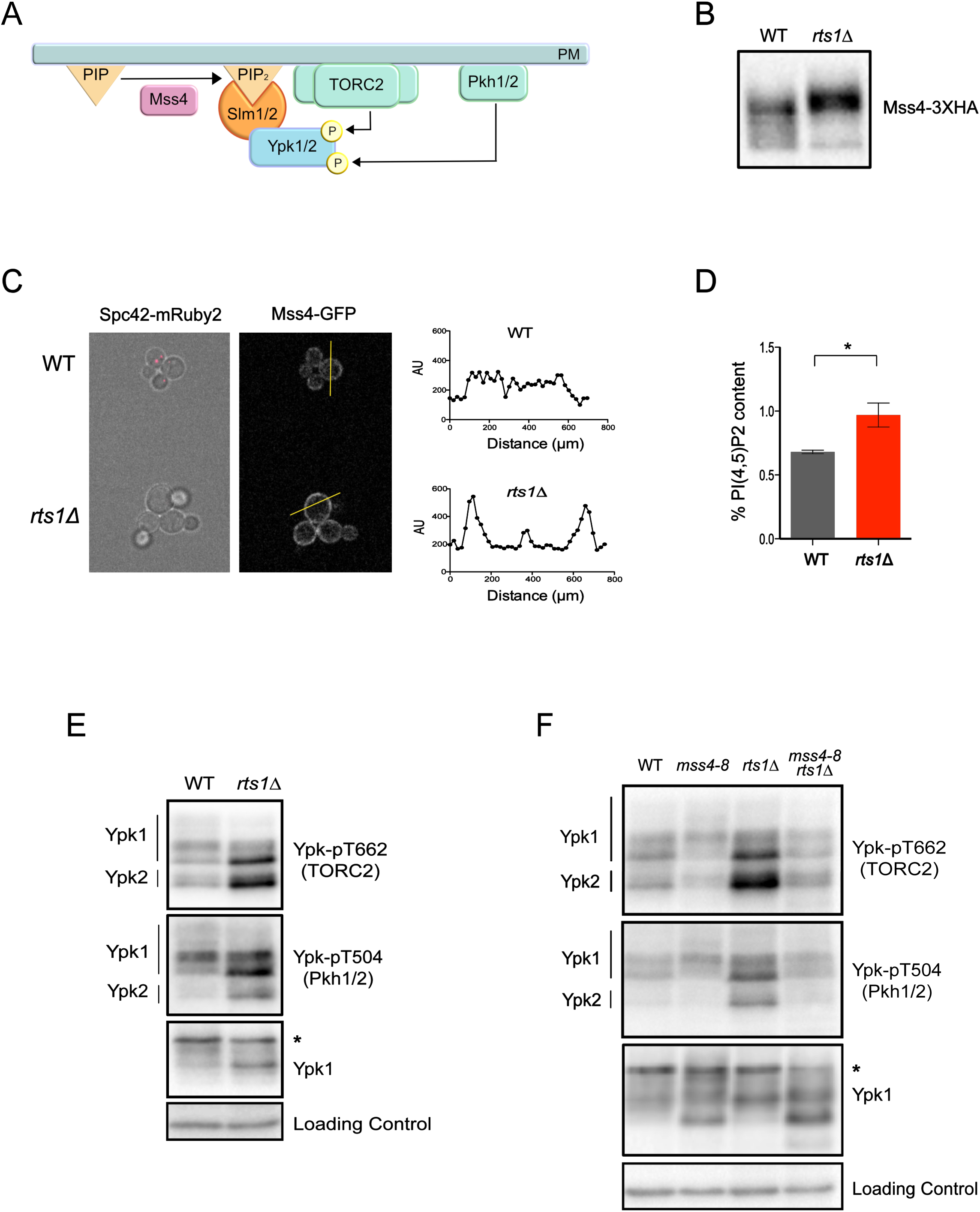
PP2A^Rts1^ controls the TORC2 signaling network via the PI(4)P kinase Mss4. **(A)** A summary of signaling events in the TORC2 network. **(B)** Wildtype and *rts1*Δ cells were grown to early log phase at 22°C in YPD medium. Mss4-3XHA was detected by western blotting. **(C)** *rts1*Δ and wildtype control cells containing Mss4-GFP were grown at 22°C in the same culture in complete synthetic medium containing dextrose. Wildtype controls cells were marked with Spc42-mRuby2 so they could be distinguished from *rts1*Δ cells. Cells were imaged by fluorescence microscopy in a field of view that includes both strains. Signal intensity was quantified along a line that bisected the plasma membrane. **(D)** Bar plots showing levels of PI(4,5)P_2_ in wildtype and *rts1*Δ cells grown to early log phase at 22°C in YPD medium. Error bars represent the standard deviation of the mean of two biological replicates. * denotes p= 0.025 in Student’s t-test. **(E)** Wildtype and *rts1*Δ cells were grown to early log phase at 22°C in YPD medium. Western blotting with phosphospecific antibodies was used to detect a TORC2-dependent phosphorylation site (Ypk-pT662) and a Pkh1/2-dependent site (Ypk-pT504) on Ypk1/2. Total Ypk1 was detected using an anti-Ypk1 antibody. **(F)** Cells of the indicated genotypes were grown overnight to early log phase at 22°C in YPD medium. The cultures were then shifted to 30°C for 2 hours. Western blotting with phosphospecific antibodies was used to detect a TORC2-dependent phosphorylation site on Ypk1/2 (Ypk-pT662) and a Pkh1/2-dependent site (Ypk-pT504). Total Ypk1 was detected using an anti-Ypk1 antibody. The asterisk indicates a non-specific band.

Mss4 undergoes phosphorylation that causes an electrophoretic mobility shift, which can be detected by western blotting. We found that Mss4 was hyperphosphorylated in *rts1*Δ cells, consistent with the proteome-wide mass spectrometry data (**Figure 1B**). Phosphorylation of Mss4 is thought to promote its association with the plasma membrane, although this has never been tested by direct visualization of plasma membrane localization. Here, we found the amount of Mss4-GFP recruited to the plasma membrane was increased in *rts1*Δ cells, consistent with the idea that hyperphosphorylation of Mss4 drives membrane association (**Figure 1C**). Increased association of Mss4 with the plasma membrane should drive increased production of PI(4,5)P_2_. We quantified PI(4,5)P_2_ in wildtype and *rts1*Δ cells, which confirmed that loss of PP2A^Rts1^ leads to increased levels of PI(4,5)P_2_ (**Figure 1D**). Thus, the data suggest that Mss4 is hyperactive in *rts1*Δ cells.

### PP2A^Rts1^ influences signaling in the TORC2 network

The discovery that Mss4 is hyperactive in *rts1*Δ cells suggested that it could drive increased TORC2-dependent hyperphosphorylation of Ypk1/2. To test this, we utilized a phosphospecific antibody that recognizes a TORC2-dependent site found on both Ypk1 and Ypk2 (**Figures S1A** **and** **S1B**). An antibody that recognizes Ypk1 was used to control for changes in Ypk1 levels. Both Ypk1 and Ypk2 showed increased TORC2-dependent phosphorylation in *rts1*Δ cells (**Figure 1E**). To test whether hyperphosphorylation of Ypk1/2 in *rts1*Δ cells is dependent upon Mss4, we generated a new temperature-sensitive allele of *MSS4* with a low restrictive temperature (30°C), which allowed us to inactivate Mss4 without the complication of heat shock effects (**Figure S2**). Inactivation of Mss4 caused loss of TORC2-dependent phosphorylation of Ypk1/2 and eliminated the increased TORC2-dependent phosphorylation of Ypk1/2 in *rts1*Δ cells (**Figure 1F**).

A phosphospecific antibody that recognizes the site on Ypk1/2 that is phosphorylated by Pkh1/2 revealed that *rts1*Δ also caused an Mss4-dependent increase in phosphorylation of Ypk1/2 by Pkh1/2 (**Figures 1E, F**). Previous studies have suggested that phosphorylation of Ypk1/2 by TORC2 promotes further activating phosphorylation of Ypk1/2 by Pkh1/2 [20,22]. Together, these observations suggest that loss of PP2A^Rts1^ causes increased Mss4 activity, which drives increased phosphorylation and activation of Ypk1/2 by TORC2 and Pkh1/2.

### PP2A^Rts1^ modulates the TORC2 network in response to nutrients

PP2A^Rts1^ is required for nutrient modulation of cell size, which suggests that it relays nutrient dependent signals [18]. We therefore considered the possibility that PP2A^Rts1^ relays nutrient-dependent signals to the TORC2 network via Mss4. However, previous studies have found no evidence that TORC2 or Pkh1/2 are modulated by nutrients in budding yeast or vertebrates. To investigate further, we first tested whether Mss4 responds to nutrient dependent signals, and whether PP2A^Rts1^ is required for the response. Wildtype cells and *rts1*Δ cells were shifted from rich carbon (2% glucose) to poor carbon (2% glycerol/ethanol) and Mss4 phosphorylation was assayed by western blotting. Mss4 underwent rapid dephosphorylation in response to poor carbon that was partially dependent upon PP2A^Rts1^ (**Figure 2A**). Purified PP2A^Rts1^ was not capable of dephosphorylating Mss4 in vitro, which suggests that it controls Mss4 indirectly, or that it requires additional factors to act on Mss4 (not shown).

**Figure 2:**
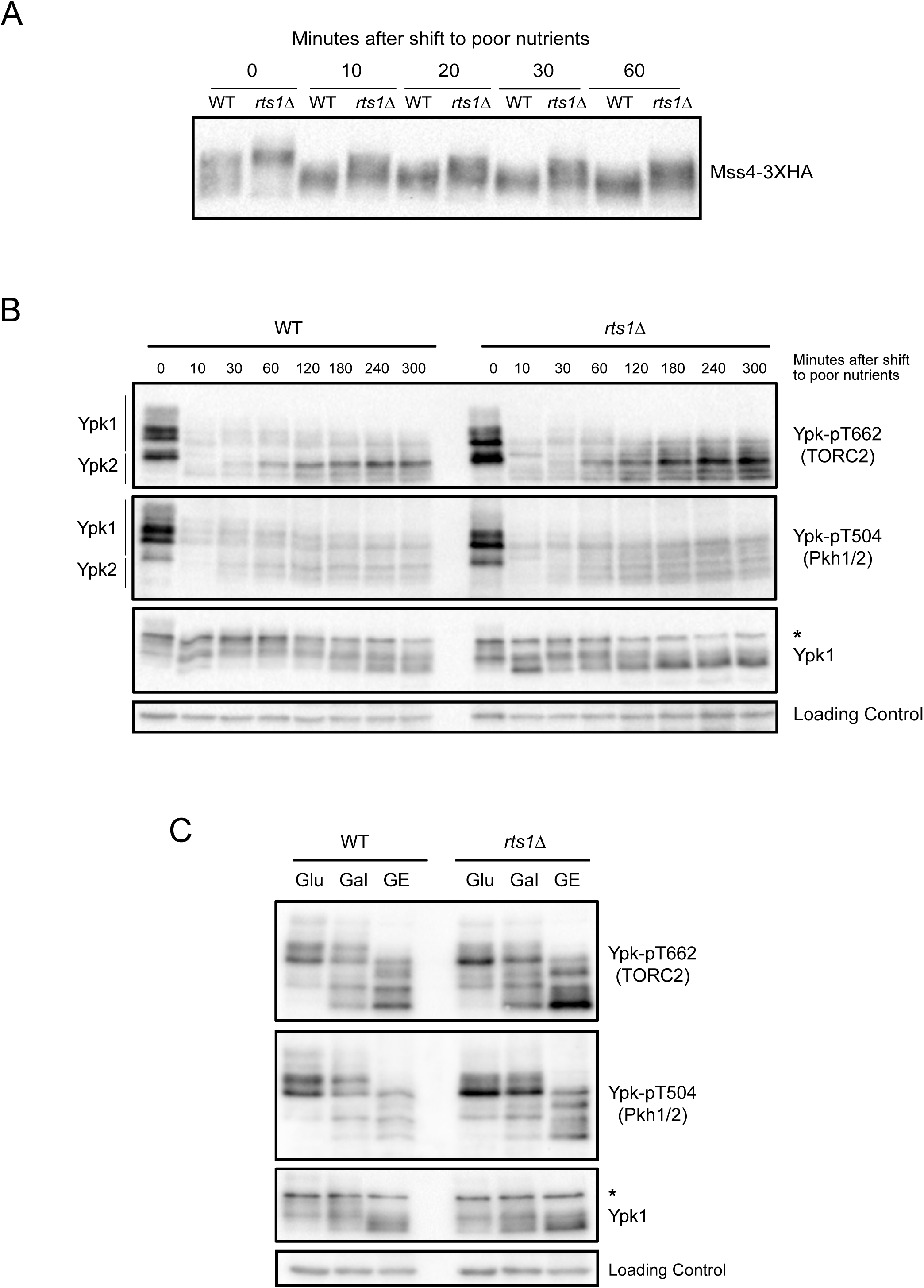
The TORC2 network is modulated by nutrients. **(A)** Wildtype and *rts1*Δ cells were grown to early log phase at 22°C in YPD medium. Cells were washed into YPG/E medium and the behavior of Mss4-3XHA was assayed by western blotting at the indicated times. **(B)** Wildtype and *rts1*Δ cells were shifted from rich to poor carbon as in (A). Ypk-pT662, Ypk-pT504 and Ypk1 were assayed by western blot. **(C)** Wildtype and *rts1*Δ cells were grown at 22°C to early log phase in YEP media containing glucose (Glu), galactose (Gal) or glycerol/ethanol (GE). Ypk-pT662, Ypk-pT504 and Ypk1 were assayed by Western blotting. An asterisk indicates a non-specific band.

We next tested whether TORC2 or Pkh1/2 signaling are modulated in response to changes in carbon source. In wildtype cells, a shift to poor carbon caused rapid loss of TORC2-dependent phosphorylation of Ypk1/2 (**Figure 2B**). As cells adapted to the new carbon source, TORC2 activity recovered but remained below the levels observed in rich carbon. Similar behavior was observed for Pkh1/2-dependent phosphorylation of Ypk1/2. The shift to poor carbon also caused increased electrophoretic mobility of Ypk1/2 (**Figure 2B**). Electrophoretic mobility shifts in Ypk1/2 are due to a pair of redundant protein kinase paralogs called Fpk1 and Fpk2, which play poorly understood roles in controlling Ypk1/2 [23]. Thus, the data suggest that Fpk1/2 activity is reduced in poor carbon.

In *rts1*Δ cells, the acute response to poor carbon occurred normally, but TORC2-dependent phosphorylation of Ypk1/2 recovered to abnormally high levels after cells adapted to the poor carbon. Pkh1/2-dependent phosphorylation of Ypk1/2 also recovered to abnormally high levels in *rts1*Δ cells, although to a lesser extent (**Figure 2B**).

To investigate TORC2 signaling after long term adaptation to poor carbon, we grew wildtype and *rts1*Δ cells for 16 hours in carbon sources of varying quality. Glycerol/ethanol was again used as a poor carbon source, while galactose served as a carbon source of intermediate quality. Again, we observed that both TORC2 and Pkh1/2 signaling were elevated in *rts1*Δ cells under all conditions, and that poor nutrients caused large shifts in the electrophoretic mobility of Ypk1/2 (**Figure 2C**).

Together, the data show that the TORC2 network is modulated by carbon source. Moreover, PP2A^Rts1^ is required for normal modulation of the TORC2 network in response to changes in carbon source.

### The level of Ypk1/2 signaling strongly influences cell size

We next tested whether cell size defects caused by loss of PP2A^Rts1^ could be caused by misregulation of Ypk1/2. Since Ypk1/2 are essential for viability, we analyzed the effects of partial loss of function caused by *ypk1*Δ or *ypk2*Δ. *ypk1*Δ caused a large decrease in cell size (**Figure 3A**). Moreover, *ypk1*Δ was epistatic to *rts1Δ*, since *rts1*Δ failed to increase the size of *ypk1*Δ cells. Loss of Ypk2 had no effect on cell size (**Figure S3A**).

**Figure 3:**
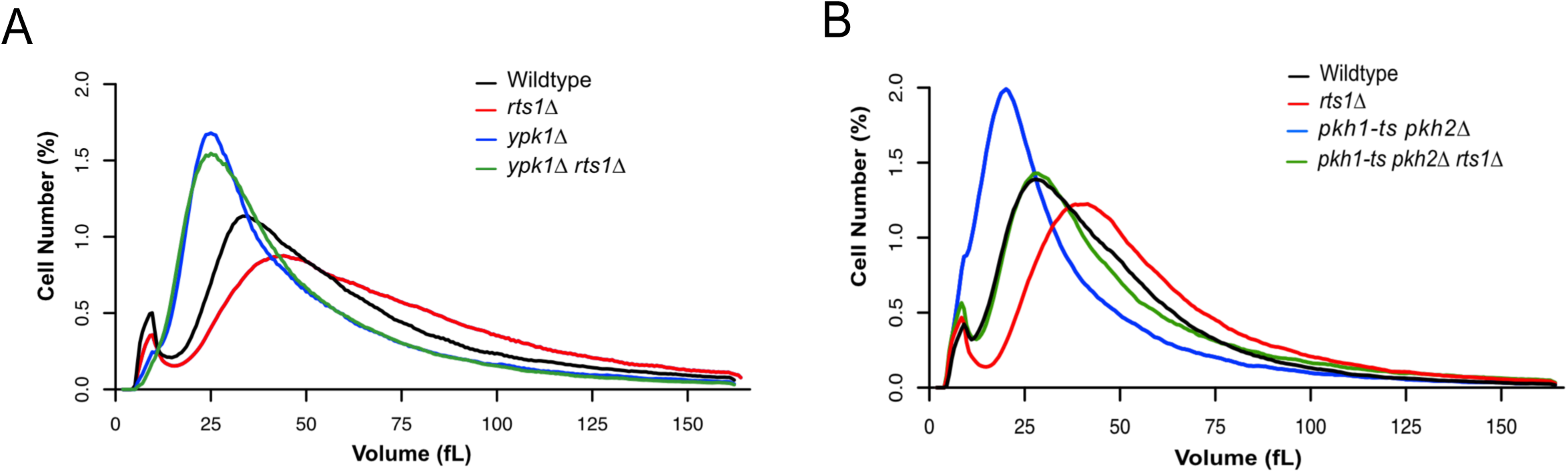
Ypk1/2 signaling strongly influences cell size. **(A,B)** Cells of the indicated genotypes were grown to log phase in YPD media at 22°C and cell size distributions were determined using a Coulter counter. Each plot is the average of 3 biological replicates. For each biological replicate, 3 technical replicates were analyzed and averaged.

We also tested whether Pkh1/2, critical upstream activators of Ypk1/2, influence cell size. To achieve partial loss of function we utilized a temperature sensitive allele of *pkh1* in a *pkh2*Δ background (*pkh1-ts pkh2*Δ) and grew cells at a semi-restrictive temperature. Partial loss of function of Pkh1/2 caused reduced cell size (**Figures 3B**). Moreover, partial loss of function of Pkh1/2 strongly reduced the size of *rts1*Δ. Overexpression of Pkh1 and Pkh2 caused increased cell size (**Figure S3B**). Previous studies in *Drosophila* and mice found that reduced PDK1 activity causes reduced cell size, while overexpression causes increased cell size, which suggests that the relationship between PDK1 signaling and cell size is conserved [24,25].

These observations demonstrate that Ypk1 activity has a strong influence on cell size. Furthermore, the data suggest that the increased size of *rts1*Δ cells is due to hyperactivity of Ypk1/2.

### Ceramides are required for normal control of cell size

An important function of Ypk1/2 is to control synthesis of sphingolipids, which are structural components of membranes and also play roles in signaling. The first step of sphingolipid synthesis is catalyzed by the enzyme serine palmitoyltransferase, which joins a long fatty acid tail to serine to create lipids called long chain bases (**Figure 4A**) [26,27]. Further processing steps use long chain bases as precursors to generate ceramides, which include two long chain fatty acid tails linked to a polar head group. The polar head group of ceramides can be further modified to generate a class of lipids called complex sphingolipids. In budding yeast, these include inositol-phosphorylceramide (IPC), mannosyl-inositol-phosphorylceramide (MIPC), and mannosyl-diinositol-phosphorylceramide (M(IP)2C). Complex sphingolipids constitute approximately 10% of total cellular lipids. In contrast, ceramides are present at approximately 0.1% of total lipids [28,29].

**Figure 4:**
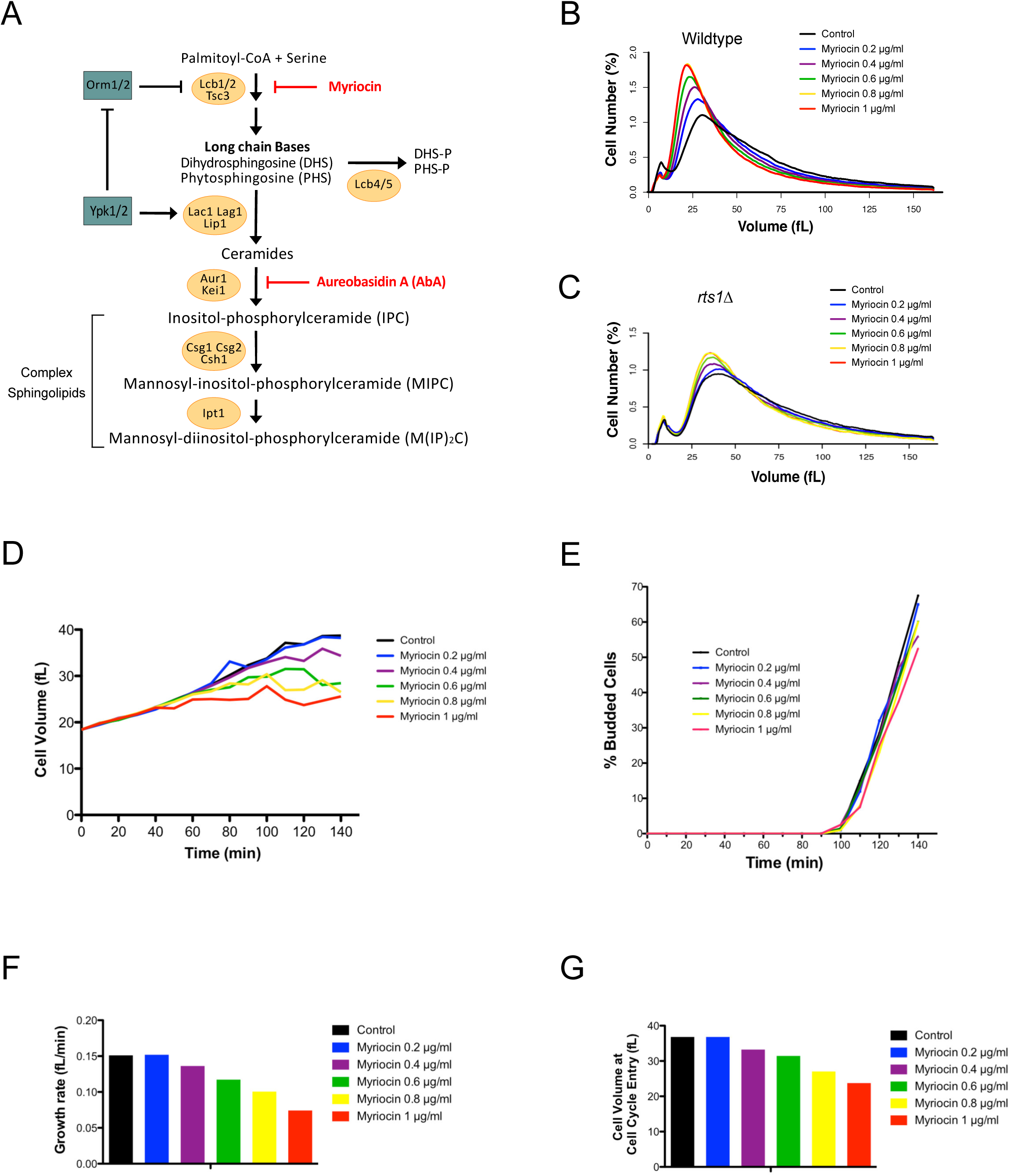
Ceramide is required for normal control of cell size. **(A)** A summary of sphingolipid synthesis pathways. Small-molecule inhibitors are indicated in red. DHS: dihydrosphingosine, PHS: phytosphingosine, DHS-P/PHS-P: dihydrosphingosine/phytosphingosine-1-phosphate. **(B,C)** Wildtype or *rts1*Δ cells were grown in YPD media at 22°C for 16 hours to early log phase in the presence of varying concentrations of myriocin. Cell size distributions were determined using a Coulter counter. **(D-G)** Cells were grown to early log phase in YPG/E medium and small unbudded cells were isolated by centrifugal elutriation. Cells were released into YPD medium in the presence of varying concentrations of myriocin at 25°C and samples were taken at 10 minute intervals. **(D)** Mean cell volume was measured with a Coulter counter and plotted as a function of time. **(E)** The percentage of budded cells was plotted as a function of time. **(F)** A plot of growth rate versus myriocin concentration. **(G)** A plot of cell size at cell cycle entry versus myriocin concentration.

Ypk1/2 promote sphingolipid synthesis by phosphorylating and inhibiting Orm1 and Orm2, a pair of redundant paralogs that bind and inhibit serine palmitoyltransferase [30,31]. Ypk1/2 also directly phosphorylate and stimulate ceramide synthase, which uses long chain base precursors to synthesize ceramides [32,33]. Ceramide synthase is encoded by redundant paralogs called LAC1 and LAG1 [34,35]. Inhibition of sphingolipid synthesis causes increased TORC2-dependent phosphorylation of Ypk1/2, as well as increased phosphorylation of the Orm1/2 proteins [30,36]. Thus, it is thought that sphingolipid synthesis is controlled by negative feedback signaling in the TORC2 network.

Since Ypk1/2 control sphingolipid synthesis, we tested whether modulation of sphingolipid synthesis could be involved in cell size defects caused by a decrease in Ypk1/2 signaling. To modulate sphingolipid synthesis, we first utilized myriocin, a small molecule inhibitor of serine palmitoyltransferase [37,38]. Cells were grown for 16 hours in sub-lethal concentrations of myriocin ranging from 0.2 μg/ml to 1 μg/ml. Previous work has shown that myriocin at these concentrations causes a dose-dependent decrease in steady state levels of long chain bases [39]. Here, we found that myriocin caused a dose-dependent decrease in cell size (**Figure 4B**). The decrease in size caused by myriocin was largely eliminated in *rts1*Δ cells, consistent with the findings that *rts1*Δ cells have increased Ypk1/2 activity and are resistant to myriocin (**Figure 4C**) [31]. The decrease in cell size was also largely eliminated by overexpression of Pkh2, which should promote increased activity of Ypk1/2. (**Figure S4A**). Myriocin also decreased the size of fission yeast, which suggests that it influences a conserved mechanism (**Figure S4B**).

To investigate further, we tested the effects of myriocin on cell growth and size at a specific stage of the cell cycle. Centrifugal elutriation was used to isolate small unbudded cells in G1 phase. Before elutriation, cells were grown in media containing poor carbon to obtain very small unbudded cells. After elutriation, cells were released into media containing rich carbon, which resulted in an extensive interval of growth as cells grew to achieve the increased size of cells in rich carbon. Myriocin was added at the same range of doses that was used to analyze the size of rapidly growing cells in **Figure 4B**. Average cell size and the percentage of cells with buds were measured at 10 minute intervals during growth in G1 phase (**Figures 4D** **and** **4E**). Growth rate was measured as the increase in volume before 25% bud emergence divided by the time (**Figure 4F**). Finally, the timing of cell cycle entry in late G1 phase was defined as the time at which 25% of the cells had buds, which was used to determine cell volume at cell cycle entry (**Figure 4G**).

Control cells grew at a rate of 0.15 fL/min during G1 and initiated bud emergence at a size of 37 fL. Myriocin caused a dose-dependent decrease in growth rate, yet had no effect on the timing of bud emergence. It also caused a dose-dependent decrease in the size at which cells entered the cell cycle. The maximal dose of myriocin decreased growth rate to 0.074 fL/min and reduced cell size at cell cycle entry to 23.70 fL.

To test how sphingolipid production could influence cell cycle entry, we analyzed how myriocin affects the behavior of Cln3 and Whi5, which are required for normal control of cell size in G1 phase. Cln3 is an early G1 cyclin that activates Cdk1, which then phosphorylates and inactivates Whi5, a transcriptional repressor that blocks transcription of genes needed for cell cycle entry [40,41]. It has been proposed that growth-dependent dilution of Whi5 plays a critical role in triggering cell cycle entry [42]. Together, these models suggested that myriocin could induce early cell cycle entry by reducing levels of Whi5, or by increasing levels of Cln3. However, addition of the highest dose of myriocin (1 μg/ml) to cells growing in G1 phase did not cause changes in Whi5 or Cln3 protein levels (**Figure S4C**).

We next determined which products of sphingolipid synthesis influence cell size. Deletion of the genes that encode ceramide synthase (*lac1*Δ *lag1*Δ) caused a large reduction in cell size (**Figure 5A**). Blocking production of MIPC or M(IP)_2_C had no effect on cell size (**Figure S5A**). To test for a role of IPC, we used the inhibitor aureobasidin A, which blocks production of IPC. However, the results were difficult to interpret. Below 5 ng/ml, aureobasidin A had no effect on proliferation or size, whereas above 15 ng/ml it completely blocked proliferation. At intermediate concentrations, there appeared to be a sharp cutoff between lethality versus no effect. More importantly, in cultures grown in intermediate doses of aureobasidin A (12.5 ng/ml) approximately 50% of cells were inviable (**Figure S5B**). Thus, in contrast to myriocin, aureobasidin A could not be used to observe the effects of modulating levels of IPC. The lethality of aureobasidin A is thought to be caused by large increase in ceramide levels [43].

**Figure 5:**
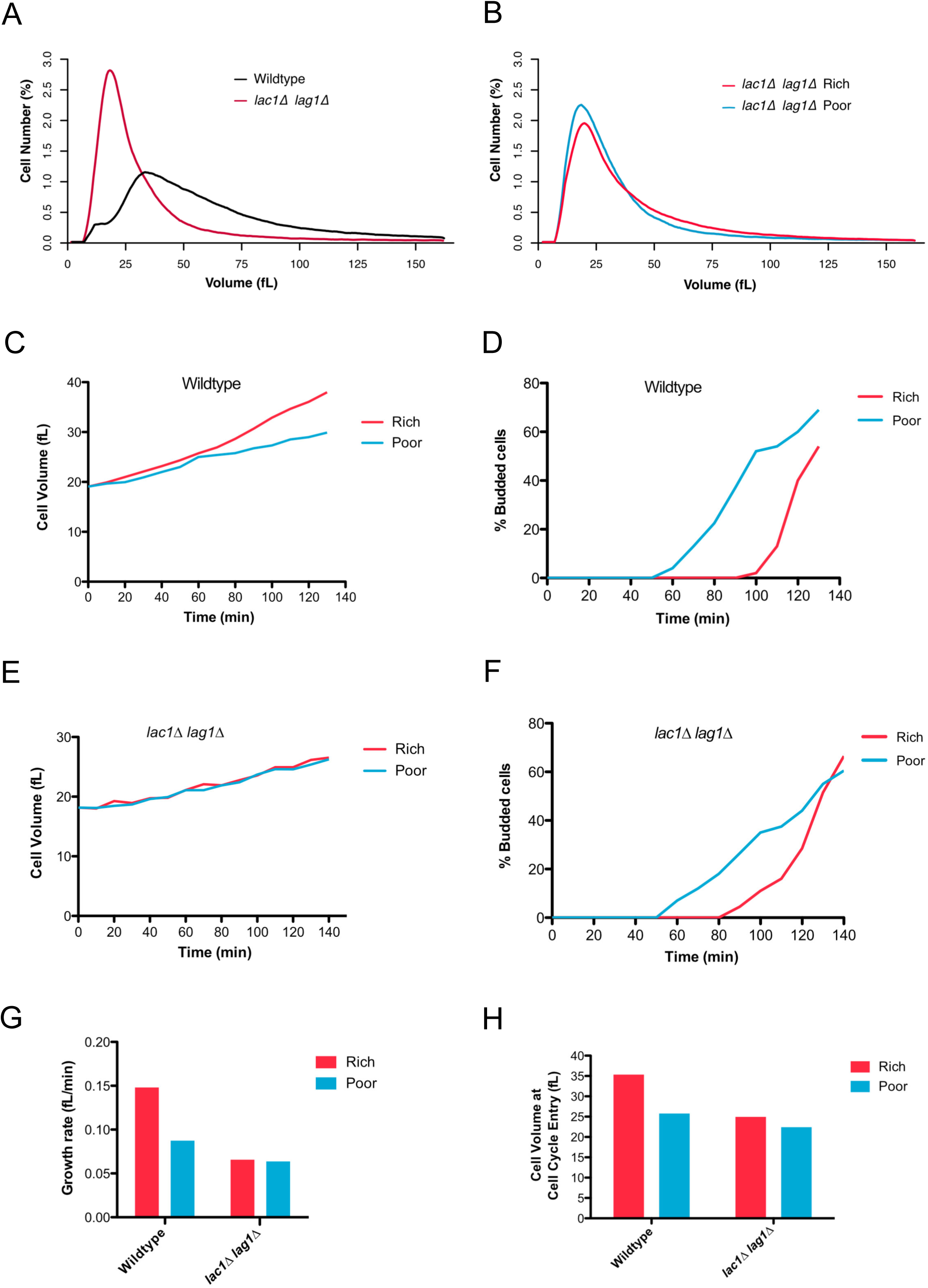
Ceramides are required for nutrient modulation of growth rate and cell size. **(A)** Wildtype and *lac1Δ lag1*Δ cells were grown to log phase in YPD medium at 22°C. Cell size distributions were determined using a Coulter counter. **(B)** *lac1Δ lag1*Δ cells were grown to log phase in YPD (Rich) or YPG/E (poor) medium for 16 hours at 22°C. Cell size distributions were determined using a Coulter counter. **(C-H)** Wildtype or *lac1Δ lag1*Δ cells were grown to log phase in YPG/E medium. Small unbudded cells in G1 were isolated by centrifugal elutriation and released into either YPD media (Rich) or YPG/E medium (Poor) at 25°C. **(C,E)** Mean cell volume was analyzed using a Coulter counter and plotted as a function of time. **(D,F)** The percentage of budded cells was plotted as a function of time. **(G)** A plot of growth rate versus carbon source. **(H)** A plot of cell volume at cell cycle entry versus carbon source.

### Ceramides are required for nutrient modulation of growth rate and cell size

PP2A^Rts1^ is required for nutrient modulation of cell size, as well as for normal modulation of signals that control ceramide synthesis. We therefore hypothesized that nutrient modulation of cell size is dependent upon modulation of ceramide synthesis. Cells that lack ceramide synthase failed to increase their size in response to rich nutrients, indicating a failure in nutrient modulation of cell size (**Figure 5B**). To investigate further, we compared growth of wildtype and *lac1*Δ *lag1*Δ cells during G1 phase in rich or poor carbon. First, small unbudded wildtype cells growing in poor carbon were isolated by centrifugal elutriation. After elutriation, half of the culture was shifted to rich carbon. The cells that remained in poor carbon grew at a rate of 0.085 fL/min and initiated bud emergence at 25.95 fL(**Figures 5C**, **5D** **and** **5G**). The cells shifted to rich carbon increased their growth rate to 0.127 fL/minute and initiated bud emergence at 35.60 fL. Bud emergence in rich carbon was delayed 40 minutes, presumably to allow cells to reach the new threshold amount of growth required for cell cycle entry in rich carbon.

In contrast, *lac1*Δ *lag1*Δ cells failed to increase their growth rate in rich carbon (**Figure 5E** **and** **5G**). Moreover, the delay in bud emergence was reduced nearly two-fold and they initiated cell cycle entry at nearly identical small sizes in rich and poor carbon (**Figures 5F** and **5H**). Thus, loss of ceramide synthase caused a nearly complete failure in modulation of growth rate and cell size in response to changes in carbon source.

### Ceramides are required for negative feedback signaling in the TORC2 network

Previous work found that signaling in the TORC2 network is influenced by negative feedback [30,36,39]. Thus, inhibition of sphingolipid synthesis triggers increased TORC2 activity. However, the mechanisms by which feedback signals are relayed are unknown. Since PP2A^Rts1^ and Mss4 influence the level of signaling in the TORC2 network, we hypothesized that they play roles in relaying feedback signals. To investigate, we tested whether Mss4 responds to sphingolipid-dependent signals. To do this, we took advantage of the fact that exogenously added phytosphingosine is taken up by cells and utilized in sphingolipid synthesis pathways. We first tested a range of phytosphingosine concentrations for effects on Mss4 phosphorylation. Addition of phytosphingosine at concentrations above 5 μM caused dephosphorylation of Mss4 (**Figure 6A**). In a time course, addition of 20 μM phytosphingosine triggered dephosphorylation of Mss4 within 10 minutes (**Figure 6B**). The strongest effects of phytosphingosine were transient, which suggests that the active lipids generated from phytosphingosine are short-lived, or that the signaling network adapts to elevated levels of phytosphingosine. Dephosphorylation of Mss4 in response to phytosphingosine was strongly attenuated in *rts1*Δ cells (**Figure 6B**).

**Figure 6:**
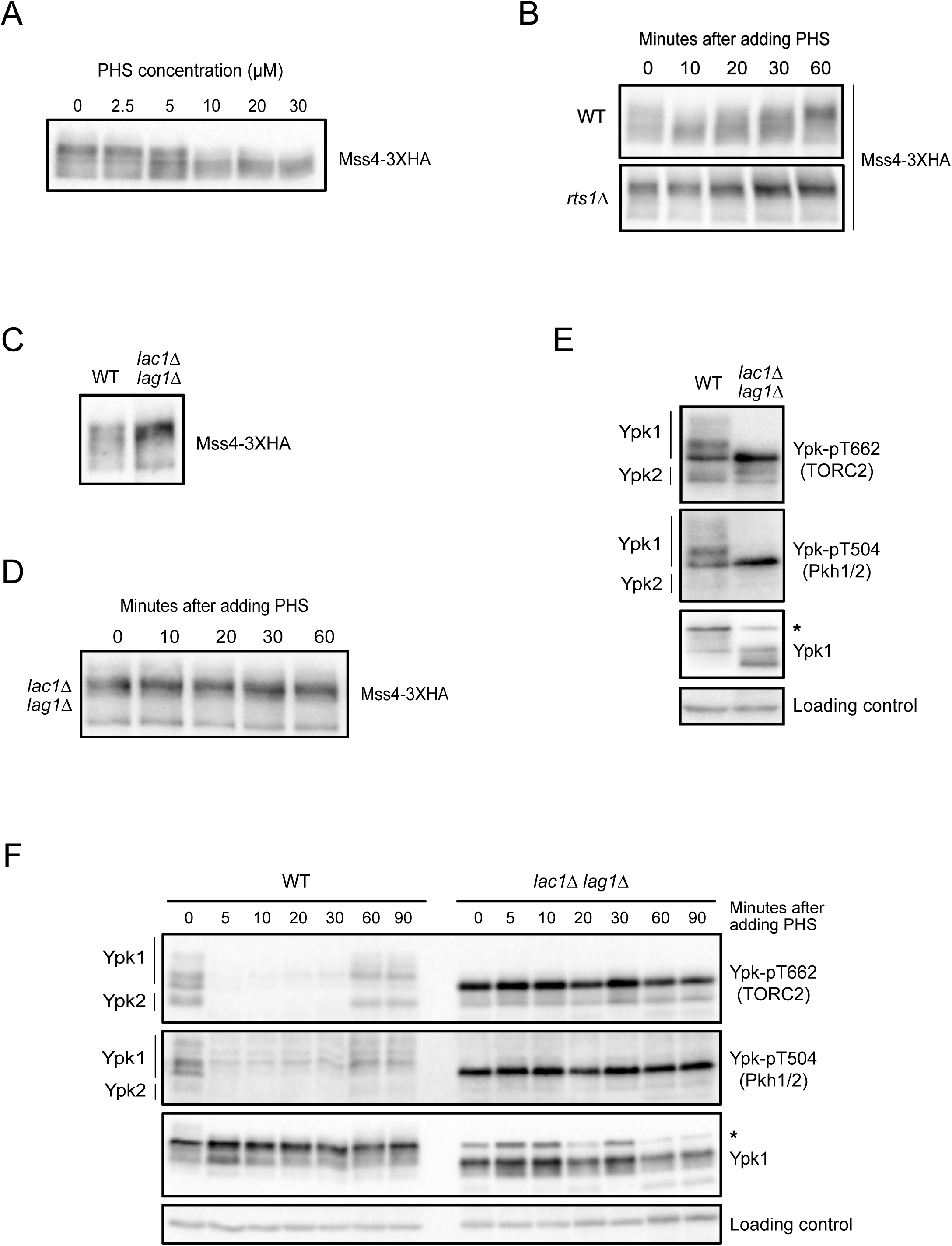
Ceramides are required for negative feedback in the TORC2 network. **(A)** Cells were grown to early log phase in YPD medium. Varying concentrations of phytosphingosine (PHS) were added and cells were incubated at 25°C. Samples were taken 15 minutes after addition of phytosphingosine and Mss4-3XHA was detected by western blot. **(B)** Wildtype and *rts1*Δ cells were grown to early log phase in YPD. 20 μM phytosphingosine (PHS) was added to each culture followed by incubation at 25°C. Samples were taken at the indicated times and Mss4-3XHA was detected by western blot. **(C)** Wildtype and *lac1*Δ *lag1*Δ cells were grown to early log phase in YPD at 22°C. Mss4-3XHA was detected by western blot. **(D)** *lac1*Δ *lag1*Δ cells were grown to early log phase in YPD at 25°C. Samples were taken at the indicated times after addition of 20 μM phytosphingosine (PHS) and Mss4-3XHA was detected by western blot. **(E)** Wildtype and *lac1*Δ *lag1*Δ cells were grown to early log phase in YPD at 22°C. Ypk-pT662, Ypk-pT504 and Ypk1 were assayed by western blot. **(F)** Wildtype and *lac1*Δ *lag1*Δ cells were grown to early log phase in YPD at 22°C. Samples were taken at the indicated times after addition of 20 μM phytosphingosine (PHS) and Ypk-pT662, Ypk-pT504 and Ypk1 were assayed by western blot. For panels E and F, an asterisk indicates a non-specific band.

We next tested whether phytosphingosine must be further modified to influence Mss4 phosphorylation. Phytosphingosine can be phosphorylated to produce sphingosine-1-phosphate, or it can be used to synthesize ceramides. Blocking phosphorylation of phytosphingosine had no effect on Mss4 phosphorylation in response to added phytosphingosine (**Figure S6A**). In contrast, Mss4 was hyperphosphorylated in *lac1*Δ *lag1*Δ cells, and dephosphorylation of Mss4 in response to exogenous phytosphingosine failed to occur (**Figure 6C,D**).

The discovery that Mss4 is hyperphosphorylated in *lac1*Δ *lag1*Δ cells predicted that TORC2 network should be hyperactive. Consistent with this, we observed increased phosphorylation of Ypk1/2 by both TORC2 and Pkh1/2 in *lac1*Δ *lag1*Δ cells (**Figure 6E**). In addition, we found that negative feedback signaling could be triggered by addition of exogenous phytosphingosine. Thus, addition of phytosphingosine to wildtype cells triggered rapid loss of TORC2 and Pkh1/2-dependent phosphorylation of Ypk1/2 (**Figure 6F**). In contrast, addition of phytosphingosine to *lac1*Δ *lag1*Δ cells failed to trigger negative feedback (**Figure 6F**).

We used aureobasidin A to test whether ceramides must be converted to complex sphingolipids to influence TORC2 signaling. To do this, we pre-incubated cells with aureobasidin A for 30 minutes to block production of IPC. This triggered a slight increase in phosphorylation of Ypk1/2 by both TORC2 and Pkh1/2, as previously reported [36]. However, aureobasidin A failed to block negative feedback signaling in response to added phytosphingosine (**Figure S6B**). Aureobasidin A also failed to block dephosphorylation of Mss4 in response to exogenous phytosphingosine (**Figure S6C**). The reason that aureobasidin A triggers increased activity of TORC2 and Pkh1/2 is unknown, although one possibility is that it has off target effects.

Together, these observations suggest that ceramides relay negative feedback signals, and that negative feedback works, at least in part, by ceramide-dependent control of PP2A^Rts1^. The data further suggest that ceramide-dependent signals stimulate PP2A^Rts1^ to dephosphorylate Mss4.

### Ceramides are required for nutrient modulation of the TORC2 signaling network

Since ceramides play a role in modulating the TORC2 network, we hypothesized that they are required for nutrient modulation of TORC2 signaling. Dephosphorylation of Mss4 in response to poor carbon was strongly reduced in *lac1*Δ *lag1*Δ cells (**Figure 7A**). In addition, the dramatic decrease in TORC2 and Pkh1/2 signaling to Ypk1/2 also failed to occur (**Figure 7B**). Thus, ceramide-dependent signals play an essential role in modulation of the TORC2 network in response to carbon source.

**Figure 7:**
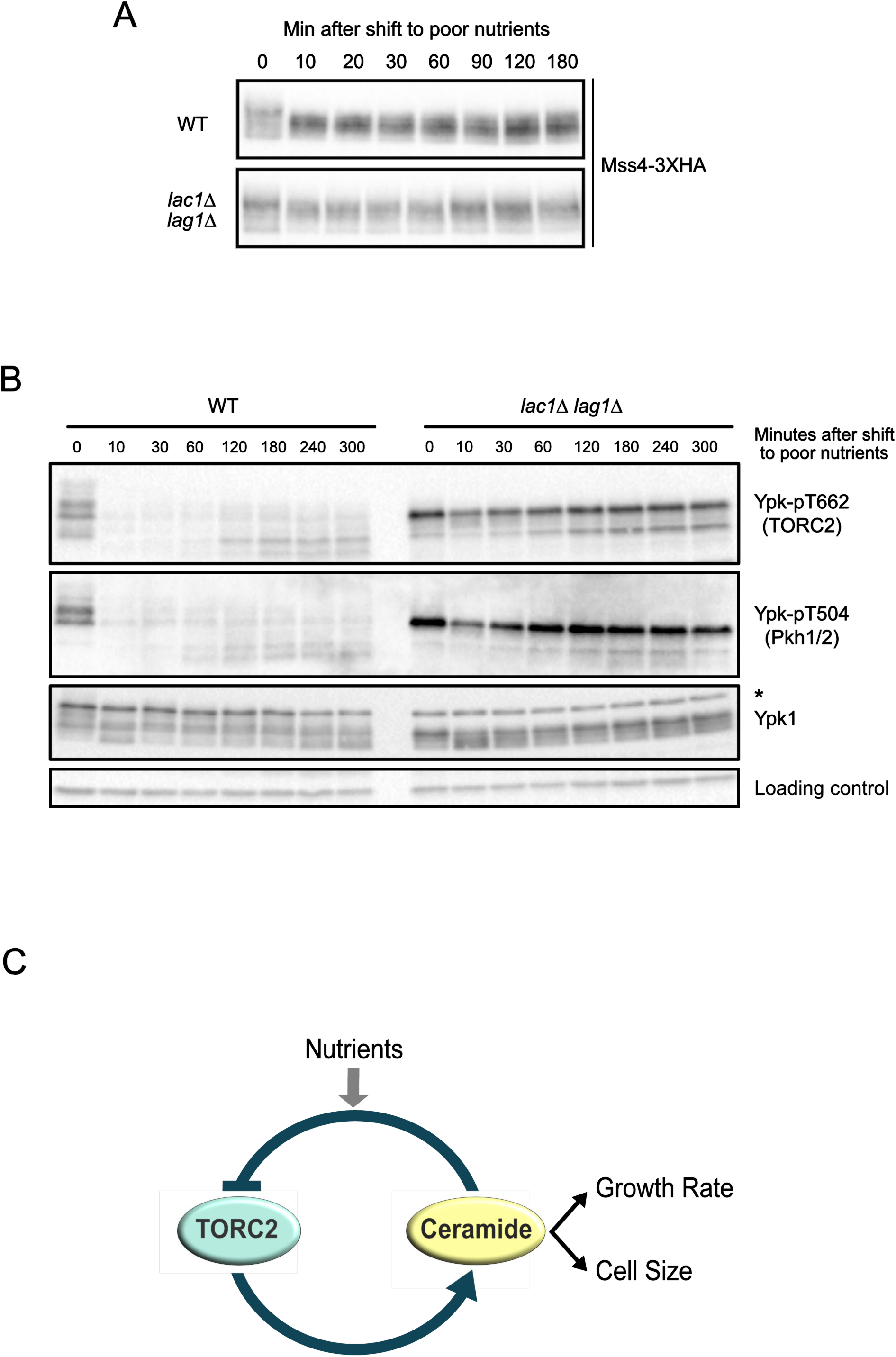
Ceramides are required for nutrient modulation of the TORC2 signaling network (A-B) Wildtype and *lac1*Δ *lag1*Δ cells were grown to early log phase in YPD at 22°C. After washing into YPG/E, cells were incubated at 25°C and samples were taken at the indicated times. Mss4-3XHA was detected by western blot. Ypk-pT662, Ypk-pT504 and Ypk1 were assayed by western blot. An asterisk indicates a non-specific band. **(C)** A model for how nutrient modulation of the TORC2 could control cell growth and size.

## Discussion

### PP2A^Rts1^ modulates TORC2 signaling in response to carbon source

It has been well-established that TORC1 is modulated by nutrients, but there has been no evidence for modulation of TORC2 by nutrients in budding yeast or vertebrates. Here, we discovered that a shift from rich to poor carbon triggers an immediate decrease in TORC2-dependent phosphorylation of Ypk1/2. Poor carbon also triggers an immediate decrease in Pkh1/2 signaling to Ypk1/2. As cells adjust to the new carbon source over several hours, TORC2 signaling recovers, but is strongly reduced relative to TORC2 signaling in rich carbon.

Diverse observations demonstrate that PP2A^Rts1^ is required for normal signaling in the TORC2 network. Loss of PP2A^Rts1^ causes hyperphosphorylation of Mss4, increased recruitment of Mss4 to the plasma membrane, and increased levels of PI(4,5)P_2_. As a consequence, TORC2 is hyperactive in *rts1*Δ cells, leading to increased phosphorylation of Ypk1/2. Pkh1/2, the other critical activators of Ypk1/2, are also hyperactive in *rts1*Δ cells. Since TORC2 phosphorylation of Ypk1/2 promotes phosphorylation of Ypk1/2 by Pkh1/2, the increased activity of Pkh1/2 in *rts1*Δ cells may be an indirect consequence of increased TORC2 activity, although it is not possible to rule out more direct effects on Pkh1/2 activity [20].

We further discovered that PP2A^Rts1^ is required for normal modulation of the TORC2 network in response to carbon source. In wildtype cells, a shift from rich to poor carbon triggers dephosphorylation of Mss4, as well as decreased activity of the TORC2 network. Both of these events fail to occur normally in *rts1*Δ cells. The data suggest a model in which PP2A^Rts1^ plays an important role in relaying information regarding carbon source to Mss4, which could modulate the level of TORC2 signaling to match the quality of the carbon source. However, it is clear that there are also PP2A^Rts1^-independent mechanisms for modulation of the TORC2 network. For example, the immediate decrease in TORC2 signaling that occurs in response to a shift to poor carbon is independent of PP2A^Rts1^. This acute PP2A^Rts1^-independent response to poor carbon could function to shut down growth as cells retool for utilization of a poor carbon source, while the long term PP2A^Rts1^-dependent response could reflect a regulated decrease in TORC2 signaling to ensure that rates of growth processes controlled by TORC2 are matched to the poor carbon source.

### PP2A^Rts1^ relays ceramide-dependent signals in the TORC2 network

Previous studies found that TORC2 signaling is strongly influenced by a negative feedback loop in which the TORC2 network promotes production of sphingolipids, while sphingolipids inhibit the TORC2 network [30,36]. However, it was unknown how feedback signals are relayed. Here, ceramides emerge as the most likely mediator of negative feedback. Inactivation of ceramide synthase causes hyperactivity of the TORC2 network. Furthermore, addition of exogenous phytosphingosine, a long chain base precursor for synthesis of ceramides, fails to trigger negative feedback in cells that lack ceramide synthase, which suggests that it must be converted to ceramide to influence TORC2 signaling. Conversion of ceramides to complex sphingolipids did not appear to be required for negative feedback signals.

Several observations suggest that ceramide-dependent feedback signals are relayed by PP2A^Rts1^. For example, addition of exogenous phytosphingosine causes PP2A^Rts1^-dependent dephosphorylation of Mss4, but fails to drive dephosphorylation of Mss4 in cells that can not make ceramide. Blocking conversion of ceramides to complex sphingolipids had no effect on the ability of exogenous phytosphingosine to signal to Mss4 via PP2A^Rts1^. Together, these observations suggest that ceramide-dependent feedback signals stimulate PP2A^Rts1^, thereby inhibiting Mss4-dependent activation of TORC2. However, the data do not rule out the possibility that there are also PP2A^Rts1^-independent mechanisms of negative feedback. Again, the observation that *lac1*Δ *lag1*Δ causes stronger effects on TORC2 network signaling than *rts1*Δ argues that ceramides do not signal solely via PP2A^Rts1^. Previous studies have suggested that ceramides could directly modulate the activity of PP2A [44,45]. Alternatively, ceramides could control PP2A^Rts1^ by recruiting it to specific locations in the cell.

The data thus far indicate that PP2A^Rts1^ could play roles in relaying both nutrient-dependent and ceramide-dependent signals. A model that could explain both roles is that PP2A^Rts1^ is embedded in the negative feedback loop so that it can relay nutrient-dependent signals that set the overall level of TORC2 network signaling.

### Ceramides are required for nutrient modulation of TORC2 signaling

In addition to a role for ceramides in feedback signaling, we discovered that ceramide-dependent signals are strictly required for nutrient modulation of TORC2 network signaling. Thus, cells that lack ceramide synthesis show a failure in modulation of the TORC2 network in response to carbon source, and they have abnormally high levels of TORC2 signaling in both rich and poor carbon. Therefore, ceramides are required for mechanisms that set TORC2 signaling to a level that is appropriate for the quality of the carbon source.

The discovery that ceramides are required for both negative feedback signaling and for nutrient modulation of the TORC2 network has important implications. Previous studies suggested that negative feedback signaling in the TORC2 network could be a homeostatic mechanism that helps maintain constant levels of sphingolipids [39]. However, sphingolipid homeostasis was only detected in cells treated with very low doses of myriocin. At higher doses, myriocin caused a dose-dependent decrease in sphingolipids [39]. Another study found that the TORC2 network responds to delivery of sphingolipids to the plasma membrane, rather than to synthesis of sphingolipids at the endoplasmic reticulum [46]. Therefore, an alternative model is that feedback ensures that the level of TORC2 signaling at the plasma membrane, which is dependent upon lipid synthesis at the endoplasmic reticulum, is maintained at a constant level that is appropriate for the carbon source. In this model, ceramide-dependent feedback ensures that TORC2 signaling is maintained at a constant level, whereas nutrient-dependent signals relayed by PP2A^Rts1^ set the overall level of TORC2 signaling, and thus the level of ceramide-dependent signaling, to a level that is appropriate for the carbon source.

### Normal control of cell size is dependent upon ceramides

In previous studies we discovered that PP2A^Rts1^ is required for nutrient modulation of cell size [2,18]. Here, we found that PP2A^Rts1^ controls signaling in the TORC2 network, and that signals from the TORC2 network are required for normal control of cell size. Thus, decreased signaling in the TORC2 network causes reduced cell size. Conversely, increased signaling, as seen in *rts1*Δ cells, causes increased cell size.

We further discovered that production of ceramides, which is controlled by the TORC2 network, is required for normal control of cell size. Inhibition of sphingolipid synthesis causes a dose-dependent decrease in cell size. In addition, inactivation of ceramide synthase causes a large decrease in cell size and a failure in nutrient modulation of cell size.

Together, the data demonstrate that an output of the TORC2 network modulates cell size. Several observations point to ceramide-dependent signals as the output that most directly modulates cell size. For example, inactivation of ceramide synthase causes decreased cell size. Similarly, decreased activity of the TORC2 network, which should cause decreased production of ceramides, also causes decreased cell size. Conversely, increased activity of the network caused by *rts1*Δ or *GAL1-PKH1/2* causes increased cell size. Most importantly, TORC2, Pkh1/2 and Ypk1/2 are all hyperactive in both *lac1*Δ *lag1*Δ cells and in *rts1*Δ cells. However, *lac1*Δ *lag1*Δ cells are unusually small, whereas *rts1*Δ cells are unusually large. A key difference is that *lac1*Δ *lag1*Δ cells fail to produce ceramides, whereas *rts1*Δ cells likely produce unusually high levels of ceramides due to hyperactivity of Ypk1/2. The implication is that ceramide levels, rather than activity of signaling proteins in the TORC2 network, are most closely correlated with cell size.

Previous work found that deletion of the gene for *SCH9*, a member of the AGC kinase family that is controlled by TORC1, causes a reduction in cell size [47,48]. Sch9 controls transcription of genes involved in ribosome biogenesis, which suggested that cell size could be linked to rates of ribosome biogenesis, although the underlying mechanisms have remained unknown. A more recent study discovered that *sch9*Δ causes reduced synthesis of ceramides [49]. This observation suggests that the reduced cell size of *sch9*Δ cells could be a consequence of a reduction in ceramide levels.

### Nutrient modulation of growth rate and cell size in G1 phase is dependent upon ceramides

To better understand how carbon source and ceramide-dependent signals influence cell growth and size, we analyzed cells growing in G1 phase. Changes in carbon source cause profound effects on cell growth and size in G1 phase. Consistent with previous reports, we found that cells shifted from poor to rich carbon in early G1 phase increase their growth rate and undergo a prolonged delay in bud emergence compared to their counterparts in poor carbon. They also enter the cell cycle at a much larger size. The most simple interpretation of these observations is that a shift from poor to rich carbon triggers an immediate resetting of the threshold amount of growth required for cell cycle entry [4,50]. As a consequence, cells shifted to rich carbon undergo a prolonged delay in cell cycle entry as they are forced to undergo additional growth to reach the higher growth threshold for cell cycle entry.

Here, we discovered that ceramide-dependent signaling strongly influences cell growth and size in G1 phase. Myriocin, an inhibitor of sphingolipid synthesis, causes a dose-dependent decrease in growth rate in G1 phase, as well as a decrease in cell size at cell cycle entry, yet has no effect on the timing of cell cycle entry. Moreover, cells that lack ceramide synthase fail to increase their growth rate when shifted from poor to rich carbon. They also fail to increase the threshold amount of growth for cell cycle entry. Thus, ceramides are required for normal nutrient modulation of growth rate and cell size in G1 phase. Cells that lack ceramide synthase still underwent a delay in bud emergence when shifted from poor to rich carbon, but the delay was cut nearly in half.

Why does loss of ceramides cause a decrease in growth rate and cell size? One might imagine that decreased production of ceramides causes decreased growth rate simply because ceramides are precursors for complex sphingolipids that are building blocks for membranes. Decreased growth rate, in turn, could lead to decreased cell size. However, we found no evidence that complex sphingolipids play roles in ceramide-dependent signals that modulate cell size or growth rate. In addition, complex sphingolipids constitute only 10% of total lipids in cells growing on rich carbon, and blocking production of MIPC and M(IP)_2_C, which constitute 50% of complex sphingolipids [28,29], had no effect on growth rate or size. Thus, the data suggest that ceramide-dependent signals could play a direct role in modulating growth rate and cell size. The idea that ceramides are the key signaling molecules is appealing because ceramides are highly conserved in eukaryotic cells, whereas complex sphingolipids show significant differences between yeast and animal cells [27].

Previous work suggested a model in which cell size at cell cycle entry is controlled by the concentration of the Whi5 protein [42]. In this model, dilution of Whi5 by growth is a critical event that triggers cell cycle entry. Here, we found that inhibition of sphingolipid synthesis with myriocin causes a large reduction in cell size at cell cycle entry without any apparent effects on Whi5 protein levels. Thus, cells treated with myriocin enter the cell cycle with higher concentrations of Whi5 relative to control cells. At the least, this observation suggests that there are mechanisms that can control cell cycle entry and cell size in G1 phase that work independently of changes in Whi5 protein concentration.

### Are growth rate and cell size coordinately controlled by common signals?

What kind of model could explain the remarkable effects of ceramide levels on cell growth and size? Also, why is cell size proportional to growth rate? One possibility is that a nutrient modulated timer determines the duration of growth in G1 phase. In this model, nutrient-dependent signals would set a timer that determines the duration of growth in G1 phase, while ceramide-dependent signals would set the growth rate. This model would be consistent with the observation that myriocin modulates growth rate without changing the timing of bud emergence. It would also be consistent with the observation that *lac1*Δ *lag1*Δ cells show a failure in modulation of growth rate in response to carbon source, yet retain a partial G1 phase delay when shifted from poor to rich carbon. Finally, it would explain why cell size is proportional to growth rate. But this model is difficult to reconcile with several observations. For example, when cells are shifted from poor to rich carbon in early G1 phase, the duration of G1 phase is dramatically increased compared to cells that remain in poor carbon. In contrast, cells that have been growing continuously in rich carbon have a much shorter G1 phase than cells growing in poor carbon, which is thought to be a consequence of cells in poor carbon being born at a smaller size and therefore having to undergo more growth in G1 to reach a threshold amount of growth for cell cycle entry [2,51,52]. Thus, the duration of growth in G1 phase is most closely correlated with the initial size of the cell and its rate of growth, rather than the carbon source, which argues against nutrient-modulated timer models.

An alternative model is that ceramide-dependent signals set growth rate, as well as the threshold amount of growth required for cell cycle entry (**Figure 7**). In this model, myriocin causes decreased growth rate, but also lowers the threshold amount of growth required for cell cycle entry. As a result, cells treated with myriocin undergo cell cycle entry with the same timing, but at different sizes. Similarly, *lac1*Δ *lag1*Δ cells fail to increase their growth rate in rich carbon, and also fail to increase the threshold amount of growth required for cell cycle entry. The observation that *lac1*Δ *lag1*Δ cells still undergo a partial G1 delay when shifted from poor to rich carbon indicates that there are ceramide-independent signals that control G1 duration. Previous studies have shown that growth in G1 phase can be divided into two intervals referred to as T_1_ and T_2_ [1,52,53]. T_1_ is the interval before Whi5 nuclear exit and shows strong evidence of cell size control. Thus, the duration of T_1_ is strongly influenced by cell size and growth rate. T_2_ is the interval between Whi5 nuclear exit and bud emergence. In cells growing in rich carbon, T_2_ is independent of cell size and behaves more like a timer. It is unknown whether T_2_ is modulated by nutrients. Thus, one possibility is that growth during T_1_ is modulated by nutrients in a ceramide-dependent manner, whereas T_2_ is modulated by nutrients in a ceramide-independent manner.

The idea that signals originating in the TORC2 network coordinately control growth rate and size would explain why loss of PP2A^Rts1^ disrupts TORC2 signaling, as well as the normal relationship between growth rate and cell size [2,18]. An alternative model is that the close relationship between growth rate and size is the consequence of mechanisms that read growth rate and set cell size accordingly. However, this seems less likely because inactivation of PP2A^Rts1^ causes a decrease in growth rate, yet cells become larger, which indicates that cell size can be uncoupled from growth rate. In addition, inactivation of PP2A^Rts1^ eliminates the linear relationship between growth rate and size [2]. Together, these observations suggest the existence of a PP2A^Rts1^-dependent mechanism that controls both growth rate and cell size.

A detailed model for how PP2A^Rts1^ and TORC2 influence cell growth and size is shown in **Figure S7**. Distinguishing alternative models will require filling in a number of gaps in our understanding of the TORC2 network. For example, the data thus far point to ceramides as the output of the TORC2 network that could directly influence growth rate and size, yet little is known about the targets of ceramide signaling in yeast, and the mechanisms by which ceramides influence feedback signaling in the TORC2 network are unknown. Candidate targets of ceramides and other derivatives of sphingolipids have been identified [54], but thus far there are no biochemically defined ceramide-binding domains in yeast. Thus, an important next step will be to better define the targets of ceramide-dependent signaling in the TORC2 network. Another important goal will be to connect ceramide-dependent signals to the downstream effectors of cell size control that influence the durations of growth during each stage of the cell cycle. Finally, discovery of the signals that control PP2A^Rts1^ could provide insight into how nutrients and ceramide-dependent signals influence the level of signaling in the TORC2 network.

## Acknowledgments

We thank members of the laboratory for advice and support. We also thank Ted Powers (UC Davis) for the Ypk-pT662 phosphospecific antibody and John Tamkun (UC Santa Cruz) for 12CA5 antibody. This work was supported by National Institutes of Health grant GM053959.

## Author Contributions

R.L., M.A-G. and D.K., designed the experiments. R.L., M.A-G, K.S., M.H., M.D., and C.M. performed the experiments. R.L., M.A-G. and D.K., wrote the manuscript.

## Materials and Methods

### Yeast strains, media and plasmids

The genotypes of all strains are listed in Table S1. All *Saccharomyces cerevisiae* strains are in the W303 background *(leu2-3,112 ura3-1 can 1–100 ade2-1 his3-11,15 trp1-1 GAL+ ssd1-d2).* One-step PCR-based gene replacement was used for construction of deletions and epitope tags at the endogenous locus (Longtine, 1998; Janke et al., 2004). Cells were grown in YP medium (1% yeast extract, 2% peptone, 40 mg/L adenine) supplemented with 2% dextrose (YPD), 2% galactose (YPGal), or 2% glycerol/ethanol (YPG/E). *Schizosaccharomyces pombe* strains were grown in standard YE media with supplements. Microscopy experiments were carried out using complete synthetic media with dextrose (CSM).

Myriocin (Sigma) was dissolved in methanol to make a 500 ng/ml stock solution. Aureobasidin A (Takara Clontech) was dissolved in methanol to make a 5 mg/ml stock solution. Phytosphingosine (Avanti Polar lipids) was dissolved in 100% ethanol to make a 10 mM stock solution.

For experiments using analog-sensitive alleles, cells were grown in YPD medium without supplemental adenine. The adenine analog inhibitors 3-BrB-PP1 or 3-MOB-PP1 were added to cultures at final concentration of 40 μM and 50 μM, respectively.

To create the *mss4-8* allele, the His3MX6 marker was integrated downstream of the *MSS4* open reading frame (Longtine et al., 1998). A fragment that contains the *MSS4* kinase domain, the HisMX6 marker, and a short region downstream of MSS4 was then amplified with Taq polymerase to introduce mutations and then transformed into a wildtype strain (Primers: GATCAGAGTCTGCAACGGCAG and GTTCACCATCGGCCTCGAGC). Transformants were selected on-HIS media and further screened for temperature sensitivity at 30°C and 37°C. To verify that the temperature-sensitive phenotype was due to mutations in *MSS4*, candidates were tested for rescue by a plasmid containing the wildtype *MSS4* gene (pMH1). We selected a clone *(mss4-8)* that was unable to grow at 30°C. DNA sequencing identified 3 mutations within the kinase domain (L506P, S539L, I672V).

### Western blotting

To ensure that protein loading was normalized in time course experiments, we determined optical densities of cultures from each strain that yield equal amounts of extracted protein. This was necessary because large cells like *rts1*Δ cells scatter light differently. We found that optical densities of 0.7 for *rts1Δ* and 0.5 for the rest of the strains yielded normalized protein loadings.

To analyze proteins from cells growing in early log phase, cultures were grown overnight at 25°C to an OD_600_ of less than 0.8. After adjusting optical densities to normalize protein loading, 1.6-ml samples were collected and centrifuged at 13,000 rpm for 30 s. The supernatant was removed and 250 μl of glass beads were added before freezing in liquid nitrogen.

To analyze cells shifted from rich to poor nutrients, cultures were grown in YPD overnight at 25°C to an OD600 of less than 0.8. After adjusting optical densities to normalize protein loading, cells were washed three times with a large volume of YPG/E medium and then incubated at 30°C in YPG/E for the time course. 1.6-ml samples were collected at each time point.

Cells were lysed into 140 μl of sample buffer (65 mM Tris-HCl, pH 6.8, 3% SDS, 10% glycerol, 50 mM NaF, 100 mM glycerophosphate, 5% 2-mercaptoethanol, and bromophenol blue). PMSF was added to the sample buffer to 2 mM immediately before use. Cells were lysed in a Mini-bead-beater 16 (BioSpec) at top speed for 2 min. The samples were removed and centrifuged for 15 s at 13,000 rpm in a microfuge and placed in boiling water for 5 min. After boiling, the samples were centrifuged for 5 min at 13,000 rpm and loaded on an SDS polyacrylamide gel.

Samples were analyzed by western blotting as previously described [55]. SDS-PAGE gels were run at a constant current of 20 mA and electrophoresis was performed on gels containing 10% polyacrylamide and 0.13% bis-acrylamide. Proteins were transferred to nitrocellulose using a Trans-Blot Turbo system (Bio-Rad Laboratories). Blots were probed with primary antibody overnight at 4°C. Proteins tagged with the HA epitope were detected with the 12CA5 anti-HA monoclonal antibody (gift of John Tamkun). Rabbit anti-phospho-T662 antibody (gift of Ted Powers, University of California, Davis) was used to detect TORC2-dependent phosphorylation of YPK1/2 at a dilution of 1:20,000 in TBST (10 mM Tris-Cl, pH 7.5, 100 mM NaCl, and 0.1% Tween 20) containing 3% Milk. Pkh1/2-dependent phosphorylation of Ypk1/2 was detected using an anti-SGK phospho-specific antibody at a dilution of 1/1000 (Santa Cruz Biotechnology, catalog number sc-16744). Total Ypk1 was detected using anti-Ypk1 antibody (Santa Cruz Biotechnology, catalog number sc-12051) at a dilution of 1:2000.

All blots were probed with an HRP-conjugated donkey anti-rabbit secondary antibody (GE Healthcare, catalog number NA934V) or HRP-conjugated donkey anti-mouse antibody (GE Healthcare, catalog number NXA931) or HRP-conjugated donkey anti-goat (Santa Cruz Biotechnology, catalog number sc-2020) for 45–90 min at room temperature. Secondary antibodies were detected via chemiluminescence with Advansta ECL reagents.

### Synchronization by Centrifugal elutriation

Cells were grown overnight at 25°C in YPG/E medium to increase the fraction of very small unbudded cells. Centrifugal elutriation was performed as previously described [56,57]. In brief, cells were elutriated at 4°C at a speed of 2,800 rpm in a Beckman Coulter J6-MI centrifuge with a JE-5.0 rotor. Small unbudded cells were released into fresh YPD or YPG/E media at 25°C and samples were taken at 10-min intervals.

### Analysis of cell size distributions and bud emergence

Cell cultures were grown overnight to early log phase at 25°C. A 900 μl sample of each culture was fixed with 100 μl of 37% formaldehyde for 1 h and then washed twice with PBS + 0.04% sodium azide + 0.02% Tween-20. Cell size was measured using a Coulter counter (Channelizer Z2; Beckman Coulter) as previously described [48]. In brief, 30 μl of fixed culture was diluted in 10 ml diluent (Isoton II; Beckman Coulter) and sonicated for 3 s before cell sizing. Each plot is the average of three independent biological replicates in which three independent technical replicates were analyzed. The size of elutriated cells was measured in the same manner except that the cells were not sonicated. The percentage of budded cells was measured by counting the number of small unbudded cells over a total of ≥200 cells using a Zeiss Axioskop 2 (Carl Zeiss) and an AxioCam HRm camera with a 63×1.4 numerical aperture objective.

### Microscopy

To visualize Mss4-GFP in wild type and *rts1*Δ cells under identical conditions, we marked wild type cells with Spc42-mRuby2 so they could be cultured and visualized together with *rts1*Δ cells. Wildtype and *rts1*Δ cells containing *MSS4-GFP* were grown together to early log phase. Fields of view containing both genotypes were imaged using a Solamere spinning disk confocal system equipped with a Yokogawa CSUX-1 scan head, Nikon TE2000-E inverted stand, Hamamatsu ImageEM **x**2 camera, and Plan Apo 60x/1.4-n.a. oil objective and controlled by Micro-Manager software. Cells were imaged in 12 z-series with a step size of 0.5 μm and analyzed using Image J. Signal at the plasma membrane was quantified using Image J Cell viability was measured by trypan blue dye exclusion. 100 μl of log phase cells were added to a 100 μl trypan blue solution (0.4% in PBS). At least 300 cells were counted for each condition.

To measure cell size in *Schizosaccharomyces pombe*, cells were fixed with calcofluor (Sigma) and length at septation was calculated using Image J.

### Anionic Phospholipid Analysis

Phosphoinositides and other anionic phospholipids were measured as deacylated lipids using anionic exchange HPLC with suppressed conductivity detection. Samples were prepared from 10^7^ cells and processed for phosphoinositide enrichment, followed by deacylation [58,59].

---

**Figure S1 (Related to Figure 1):**
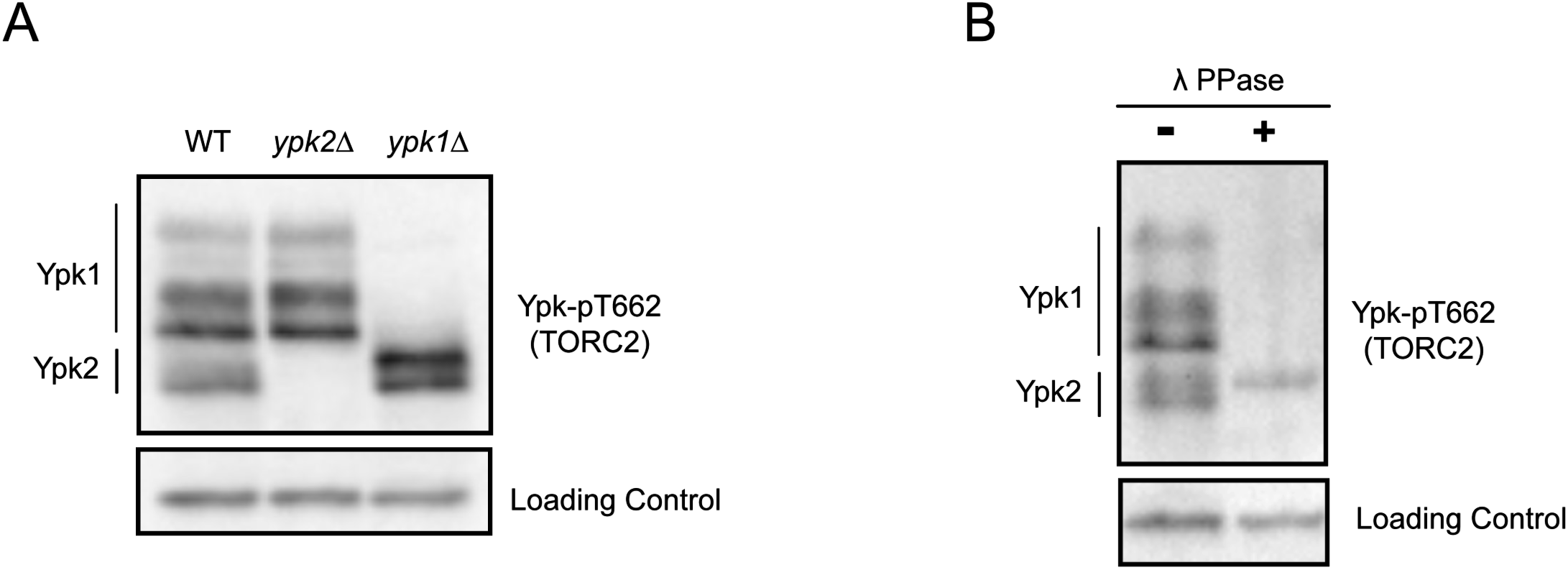
Characterization of a phosphospecific antibody that recognizes TORC2-dependent phosphorylation of Ypk1 and Ypk2. **(A)** Extracts from wildtype, *ypk1*Δ and *ypk2*Δ cells were analyzed by western blot to detect a TORC2-dependent phosphorylation site on Ypk1 and Ypk2. The antibody detects phosphorylated T662 in Ypk1 and a homologous site in Ypk2. **(B)** Extracts from wildtype cells were treated with lambda-phosphatase and Ypk-pT662 phosphorylation was assayed by western blot.

**Figure S2 (Related to Figure 1):**
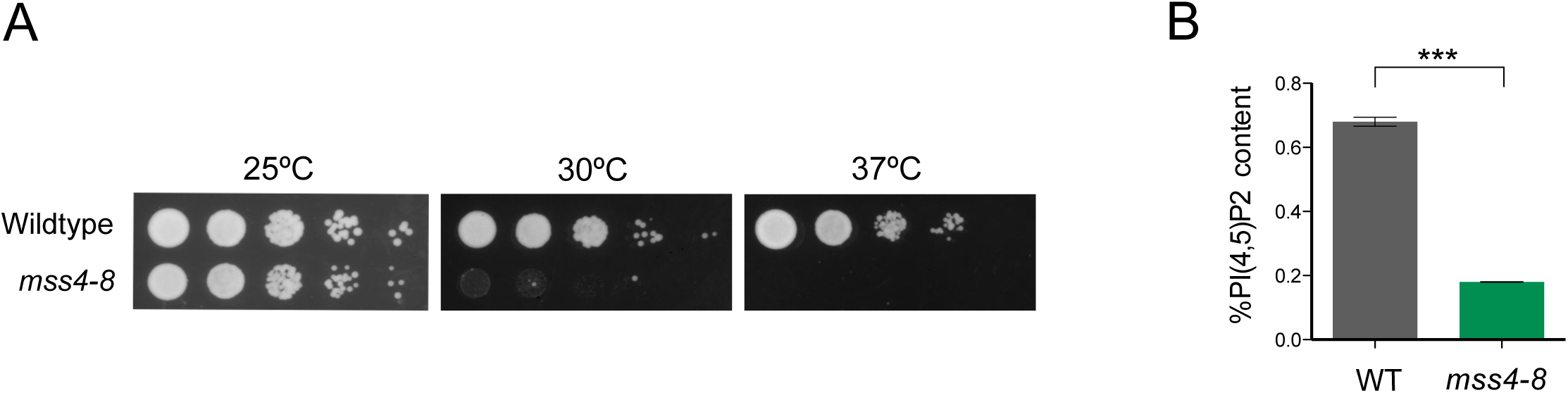
Characterization of a new temperature-sensitive allele of *MSS4*. **(A)** A series of 10-fold dilutions of cells were grown at 25°C, 30°C and 37°C on YPD. **(B)** Bar plots showing levels of PI(4,5)P_2_ in wildtype and *mss4-8* cells grown to early log phase in YPD medium at 25°C, which is a semi-restrictive temperature. Error bars represent the standard deviation of the mean of two biological replicates. *** denotes p= 0.0002 in Student’s t-test.

**Figure S3 (Related to Figure 3):**
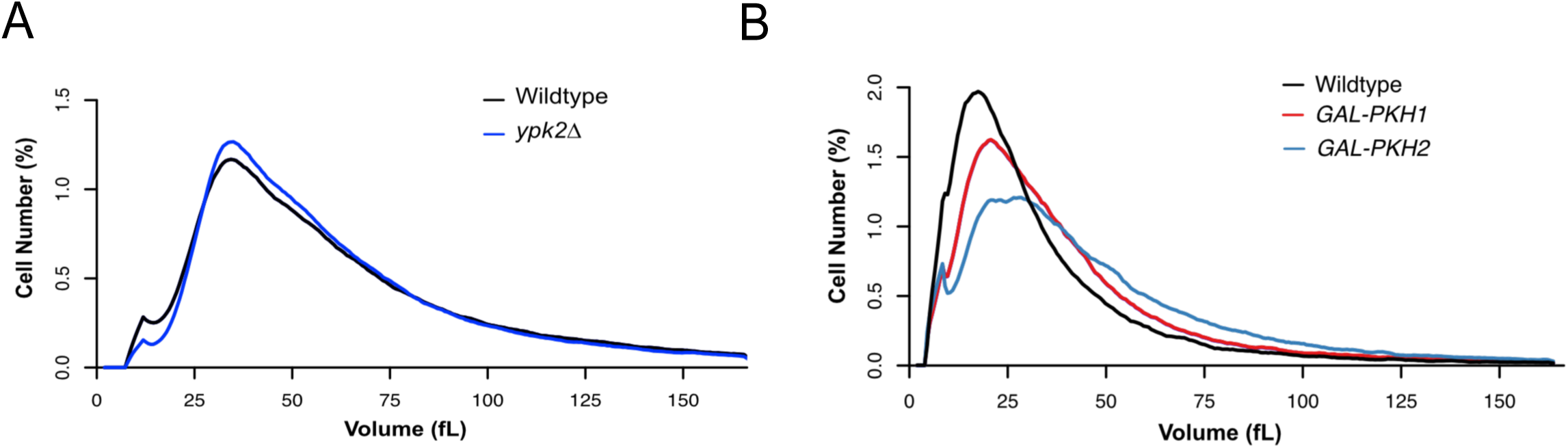
*ypk2*Δ has no effect on cell size and overexpression of *PKH1* or *PKH2* causes increased size. **(A,B)** Cells of the indicated genotypes were grown to log phase at 22°C and cell size distributions were determined using a Coulter counter. **(A)** Cells of the indicated genotypes were grown in YPD medium at 22°C. **(B)** Cells of the indicated genotypes were grown in YPGal medium at 22°C.

**Figure S4 (Related to Figure 4):**
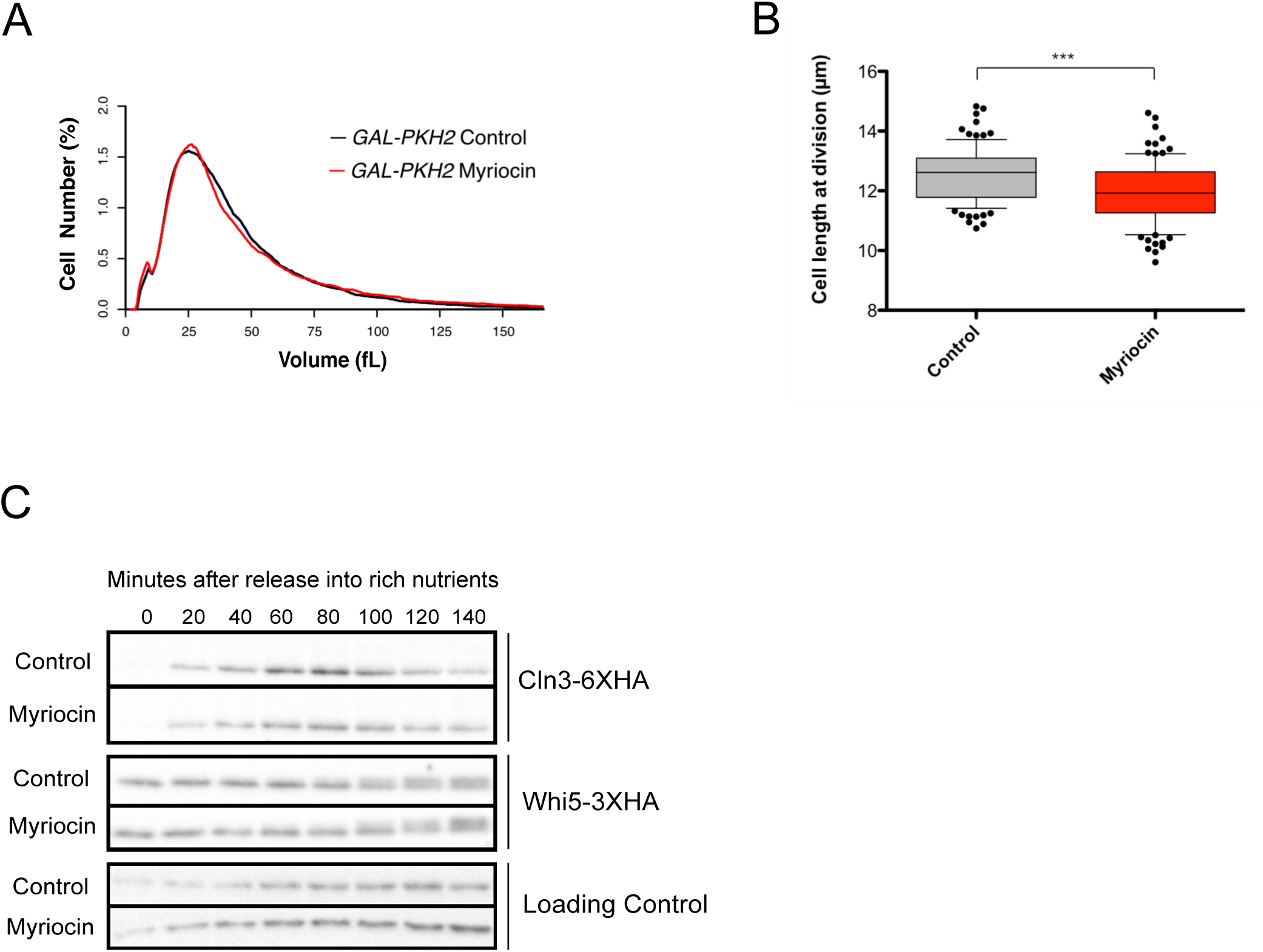
Overexpression of PKH2 suppresses effects of myriocin on cell size, myriocin causes reduced cell size in fission yeast, and myriocin does not cause changes in Whi5 or Cln3 protein levels during G1 phase. (A) *GAL1-PKH2* cells were grown in YPGal medium at 22°C for 16 hours to early log phase with or without 0.4 μg/ml myriocin. Cell size distributions were determined using a Coulter counter. **(B)** Cell length at division of wildtype fission yeast cells grown for 16 hours at 22°C in the presence or absence of 0.3 μg/ml myriocin. Boxes are delimited by the 25-75% of the data and the central lines indicate the median. Whiskers mark maximum and minimum values within a 10-90% range of the data; individual dots represent cells outside the range. *** indicates a p-value of 0.0001 in Student’s t-test **(C)** Cells were grown to early log phase in YPG/E medium. Small unbudded cells were isolated by centrifugal elutriation and were released into YPD medium with or without 1 μg/ml myriocin at 25°C. Samples were taken at 20 minute intervals and Cln3-6XHA and Whi5-3XHA were assayed by western blot. We consistently observed that cells carrying Cln3-6XHA and Whi5-3XHA budded prematurely at a reduced volume, which suggests that the tags cause a slight perturbation of function.

**Figure S5 (Related to Figure 5):**
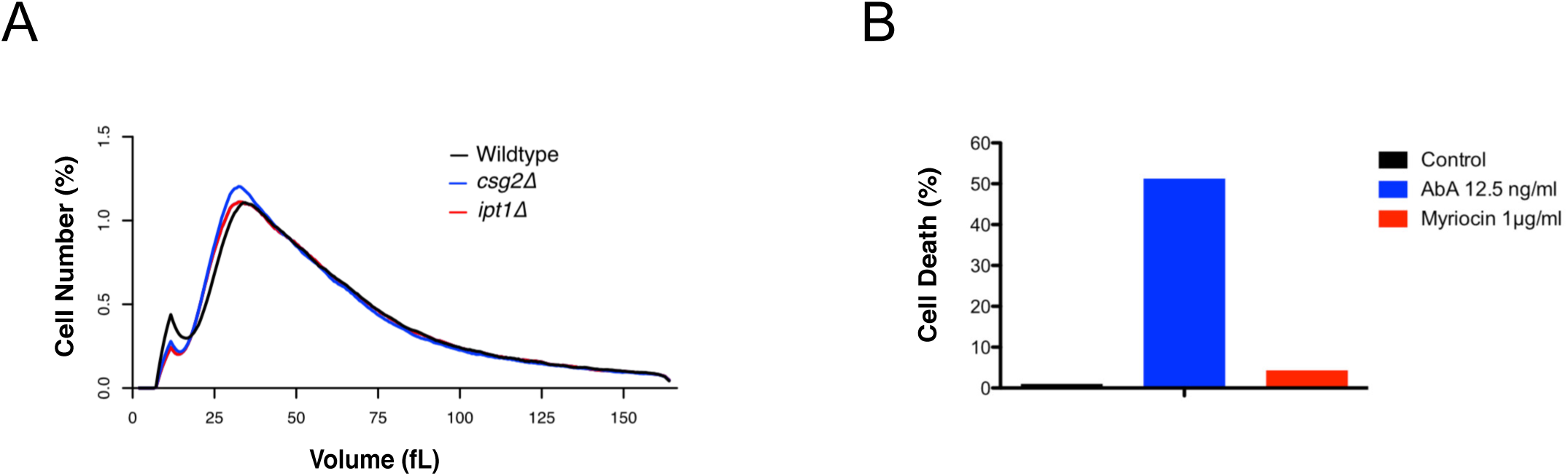
Blocking synthesis of complex sphingolipids does not cause effects on cell size. **(A)** Cells of the indicated genotypes were grown to log phase in YPD at 22°C and cell size distributions were determined using a Coulter counter. Csg2 is required for production of MIPC and Ipt1 is required for production of M(IP)2C. **(B)** Wildtype cells were grown to log phase at 22°C in YPD medium or YPD medium containing 12.5 ng/ml Aureobasidin A (AbA) or 1 μg/ml myriocin. Cell viability was determined by trypan blue exclusion assay.

**Figure S6 (Related to Figure 6):**
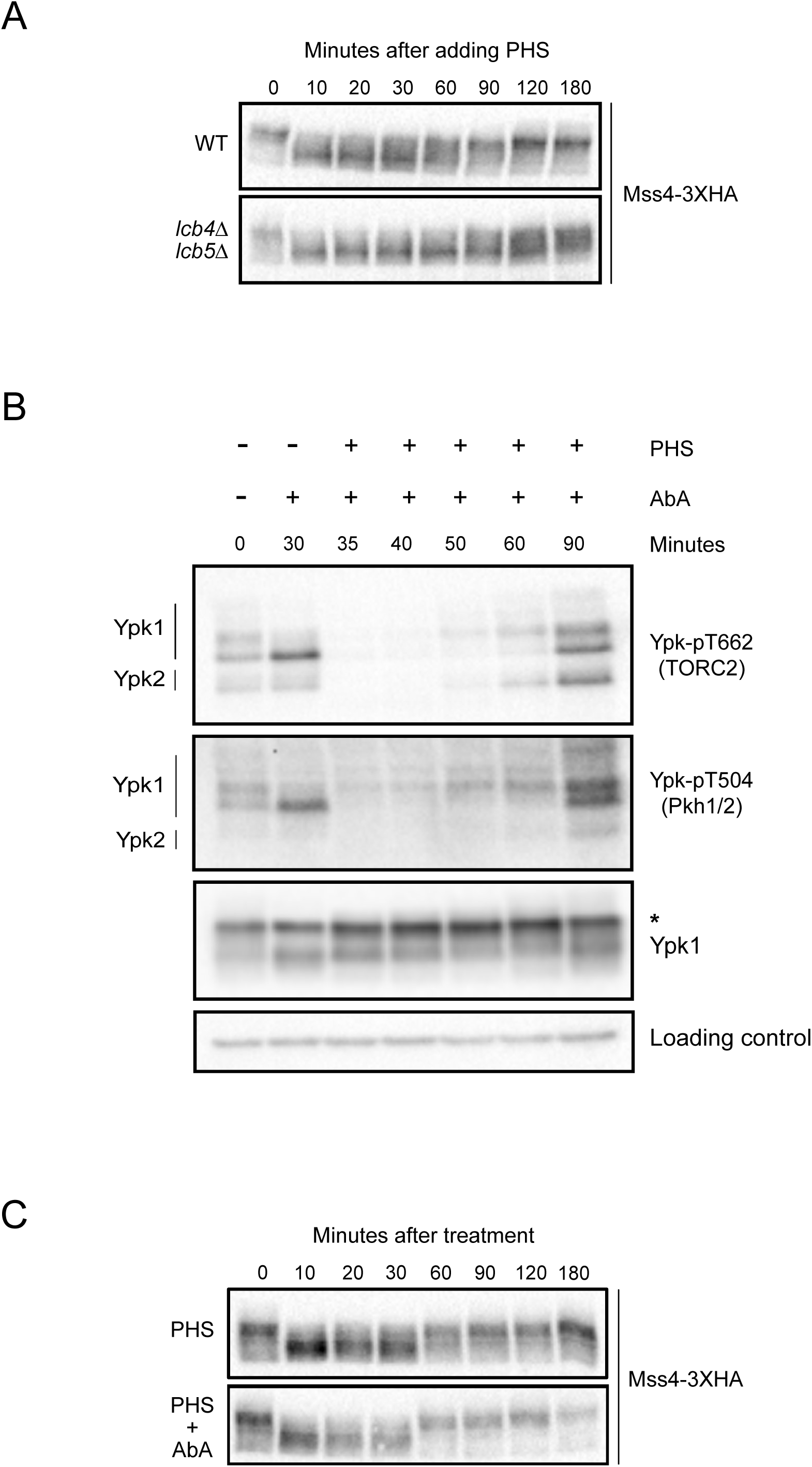
Phytosphingosine must be converted to ceramides to influence TORC2 network signaling. **(A)** Wildtype and *lcb4Δ lcb5*Δ cells were grown to early log phase at 22°C in YPD medium. 20 μM phytosphingosine (PHS) was added to each culture followed by incubation at 25°C. Samples were taken at the indicated times and Mss4-3XHA was detected by western blot. Lcb4/5 are required for phosphorylation of long chain bases. **(B)** Wildtype cells were grown to early log phase in YPD and 0.5 μg/ml Aureobasidin A (AbA) was added to the cultures followed by 30 min incubation at 25°C to ensure complete inhibition of IPC synthesis and depletion of IPC. After this time, 20 μM phytosphingosine (PHS) was added. Samples were taken at the indicated times and western blotting with phosphospecific antibodies was used to detect a TORC2-dependent phosphorylation site (Ypk-pT662) and a Pkh1/2-dependent site (Ypk-pT504) on Ypk1/2. Total Ypk1 was detected using an anti-Ypk1 antibody. **(C)** Wildtype cells were grown to early log phase in YPD medium. 20 μM phytosphingosine (PHS) or 20 μM phytosphingosine and 0.5 μg/ml Aureobasidin A (AbA) were added to the cultures, followed by incubation at 25°C. Samples were taken at the indicated times and Mss4-3XHA was detected by western blot.

**Figure S7.**
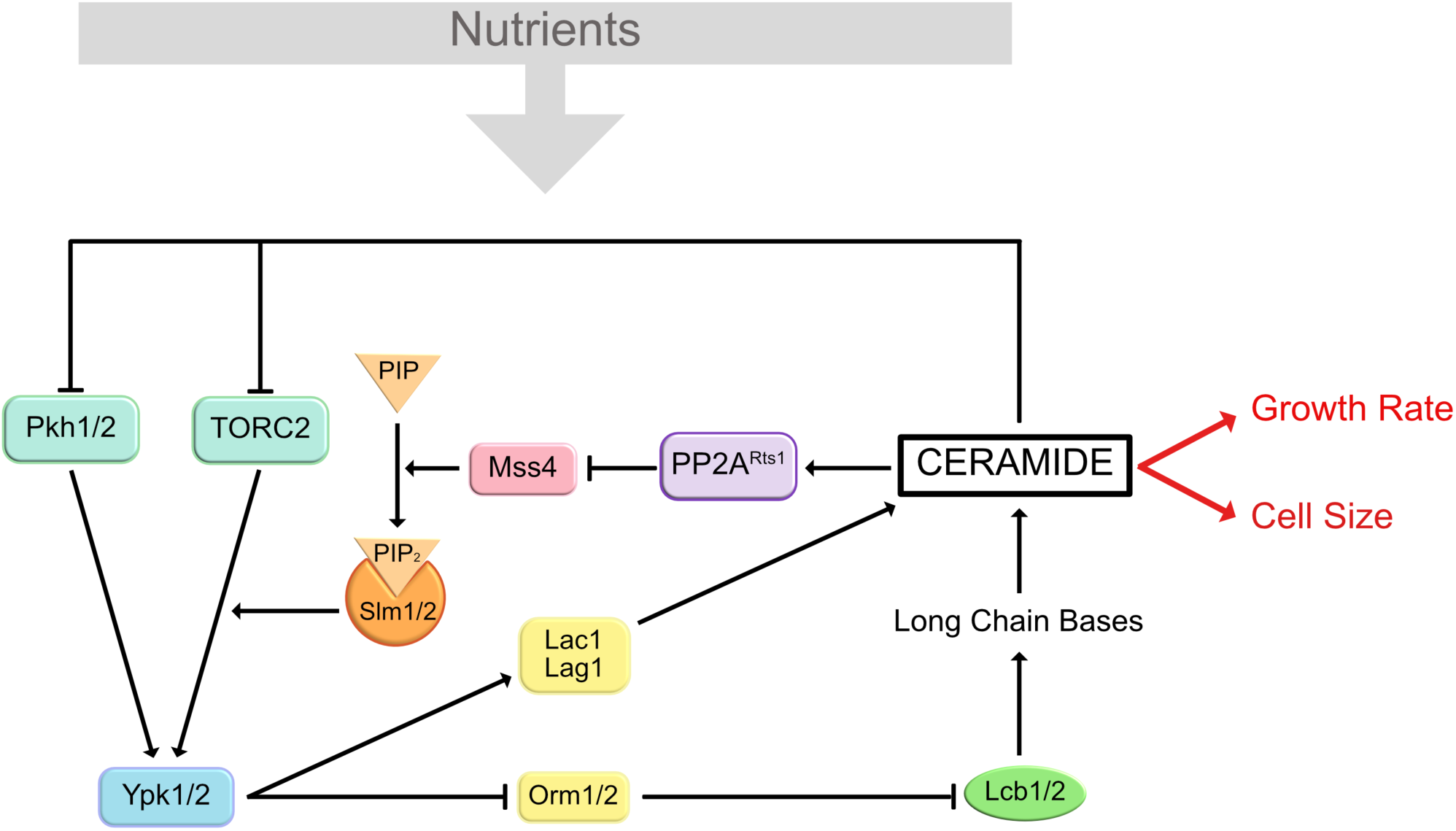
PP2A^Rts1^ and TORC2 influence cell growth and size. A model for control of cell growth and size by PP2A^Rts1^ and the TORC2 network.

**Table S1.** Strains used in this study

| Strain | MAT | Genotype | Source |
| --- | --- | --- | --- |
| DK186 | a | *bar1* | Altman and Kellogg, 1997 |
| DK647 | a | *bar1 rts1Δ::kanMX6* | Artiles et al., 2009 |
| DK2326 | a | *bar1 MSS4-3XHA::His3MX6* | This study |
| DK2342 | a | *bar1 MSS4-3XHA::His3MX6 rts1Δ::kanMX6* | This study |
| DK3157 | a | *bar1 MSS4-GFP::His3MX6 Spc42-mRuby2::KanMX6* | This study |
| DK2862 | a | *bar1 MSS4-GFP::His3MX6 rts1Δ::KanMX6* | This study |
| DK2925 | a | *bar1 mss4-8::His3MX6* | This study |
| DK2967 | a | *bar1 mss4-8::His3MX6 rts1Δ::KanMX6* | This study |
| DK3116 | a | *bar1 ypk1Δ::His3MX6* | This study |
| DK3158 | a | *bar1 ypk1Δ::His3MX6 rts1Δ::KanMX6* | This study |
| DK1523 | a | *bar1 pkh1-D398G pkh2Δ::LEU2 Cln2-3XHA::ADE2* | This study |
| DK1721 | a | *bar1 pkh1-D398G pkh2Δ::LEU2 Cln2-3XHA::ADE2 rts1Δ::His3MX6* | This study |
| DK3092 | a | *bar1 Cln3-6XHA:: His3MX6 Whi5-3XHA::hphNT1* | This study |
| DK3267 | a | *bar1 lag1Δ::kanMX6 lac1Δ:: His3MX6* | This study |
| DK3268 | a | *bar1 lag1Δ::kanMX6 lac1Δ:: His3MX6 Mss4-3XHA::TRP* | This study |
| DK2546 | a | *bar1 ypk2Δ::HIS3* | This study |
| DK2758 | a | *bar1 GAL1-3XHA-PKH2::URA* | This study |
| DK2759 | a | *bar1 GAL1-3XHA-PKH1::URA* | This study |
| DK1037 | h- | 972 (*S. pombe* wild type) | Paul Nurse |
| DK3184 | a | *bar1 csg2Δ::KanMX6* | This study |
| DK3186 | a | *bar1 ipt1Δ::KanMX6* | This study |
| DK3216 | a | *bar1 MSS4-3XHA::His3MX6 lcb4Δ::NatNT2 lcb5Δ::KanMX6* | This study |

